# Tuberculous meningitis alters the proteomic landscape of brain-derived extracellular vesicles

**DOI:** 10.64898/2025.12.08.692466

**Authors:** Blanca V. Rodriguez, Nerketa N. L. Damiba, Nicole Beaubien, Daisy Puca, D. Brian Foster, Samarjit Das, Elizabeth W. Tucker

## Abstract

Tuberculous meningitis (TB meningitis), the deadliest form of *Mycobacterium tuberculosis* infection, leads to mortality and severe neurological disability despite standard therapy. Brain injury and microglial activation are major determinants of outcome, yet the mechanisms linking infection, inflammation and neuronal injury remain poorly understood. Extracellular vesicles (EVs), key mediators of cell-to-cell communication, have been investigated in pulmonary TB but their role in TB meningitis remains unexplored. We used our young rabbit model of TB meningitis to isolate pure, intact EVs from brain tissue (i.e., brain-derived EVs) from infected and uninfected rabbits and used nanoflow cytometry, transmission electron microscopy and protein quantification to characterize the EVs. Comparative proteomic profiling was performed by liquid chromatography-tandem mass spectrometry (LC-MS/MS), followed by *in silico* pathway, cell-type and protein-protein interaction analyses using DAVID, Enrichr, and STRING databases. We found that EV isolation from fresh and frozen tissue was equivalent and demonstrated that *M. tuberculosis* infection activated EV biogenesis. Despite preserved vesicle morphology, EVs from infected brain showed a significant proteomic shift characterized by enrichment of TB host defense, microglial and immune activation, metabolic excitotoxicity, and neuronal injury. These proteome dysregulations suggest that infection reprograms brain EV cargo toward proinflammatory and metabolic stress responses while depleting neuronal and mitochondrial components. Collectively, these data demonstrate that *M. tuberculosis* infection alters the cargo and abundance of brain-derived EV, highlighting their potential as biomarkers and mediators of host-pathogen interactions in TB meningitis.

## INTRODUCTION

Tuberculous meningitis (TB meningitis) is caused by *Mycobacterium tuberculosis* and is the most severe and life-threatening form of tuberculosis (TB) that disproportionately impacts children and immunocompromised individuals. It is characterized by high rates of mortality and life-long morbidity^1,2^, with the majority of survivors left with neurological disability, including physical and neurodevelopmental deficits^2–6^. These poor outcomes occur despite appropriate antimicrobial and current standard of care host-directed therapy (corticosteroids), underscoring the need to improve our understanding of TB meningitis pathology. Brain injury is an important factor influencing outcomes in TB meningitis, with evidence from patients that elevated brain injury biomarkers (e.g., glial fibrillary acidic protein [GFAP]) predict death^7^. Glutamate excitotoxicity is also elevated, particularly among patients with seizures^8,9^. Additionally, cell, animal, and human studies demonstrate the critical role that microglia and astrocytes have in amplifying the inflammatory cascade that leads to cytokine release and neuronal injury^10–14^. Despite these insights, the precise molecular pathways leading to neuroinflammation and neuronal damage in TB meningitis remain poorly understood.

Extracellular vesicles (EVs) are crucial mediators of intercellular communication under normal conditions and pathological states like infection and inflammation. Their bioactive cargo (e.g., proteins, lipids, nucleic acids) can help maintain homeostasis or propagate neuroinflammation. EVs proinflammatory role is well-documented across several disease states, from infections like viral encephalitis to neurodegenerative diseases like Alzheimer’s disease^15,16^. Brain-derived EVs are increasingly recognized as key regulators in neurodevelopment, neurodegenerative diseases and neuroinflammation where their secretion is increased.^17,18^ To date, EV research in TB has focused on pulmonary TB, where EVs from *M. tuberculosis* have been utilized as a biomarker of active disease^19^ and EVs from host immune cells have been shown to improve bacterial clearance by activating macrophages and promoting phagosome maturation^20,21^. In contrast, the cellular origin, cargo and function of EVs from the central nervous system (CNS) in TB meningitis remain unknown. Defining the contribution of brain-derived EVs to infection could uncover novel diagnostic biomarkers and mechanistic insights into neuroinflammatory signaling and brain injury.

To address this knowledge gap, we optimized a workflow to isolate brain-derived EVs from rabbits with and without experimentally induced TB meningitis and used nanoflow cytometry (NanoFCM), transmission electron microscopy (TEM) and protein quantification to characterize the EVs. We performed the first proteomic analysis of brain-derived EVs to characterize the qualitative changes induced during infection and performed *in silico* pathway, cell-type and protein-protein interaction analyses using DAVID, Enrichr, and STRING databases to define the protein landscape. Our study provides the first evidence that *M. tuberculosis* infection reprograms brain-derived EV cargo, highlighting their potential as both biomarkers and mediators of CNS host-pathogen interactions.

## MATERIALS AND METHODS

### Animal model

We utilized New Zealand White rabbits (Envigo Global Services Inc.) for all rabbit studies. Rabbits were allowed at least two weeks to acclimatize prior to any intervention. All protocols were approved by the Johns Hopkins University Animal Care and Use (RB22M351) and Biosafety committees.

### Young rabbit model of TB meningitis

We utilized our young rabbit model of TB meningitis and healthy controls as previously described^10,11,22,23^. In brief, rabbits were bred in-house and kits were delivered spontaneously. Genotyping for sex determination was performed using a 2 mm ear sample (Transnetyx, Memphis, TN) prior to intervention. Young rabbits (kits) at postnatal day 4-8 received dexmedetomidine intramuscular (IM) and topical lidocaine to skull for analgesia and sedation prior to intraventricular injection of 20 µL of phosphate buffered saline (PBS, uninfected) or *M. tuberculosis* H37Rv inoculum from frozen stock (i.e., infected) via the bregma (intersection of the coronal and sagittal suture).

### Healthy adult rabbits

Healthy adult rabbits (female and male) used for in-house breeding were utilized to optimize EV isolation techniques prior to testing in tissue and biofluids from young rabbits due to limitations from smaller volumes.

### Brain-derived EV isolation

#### Sample collection

Young rabbits with TB meningitis (i.e., infected) and age-matched healthy controls (uninfected) were followed for three weeks unless otherwise specified and euthanized with pentobarbital overdose (120 mg/kg; VetOne, Boise, ID) for sample harvest. Three weeks were chosen due to development of symptoms and lesions on gross pathology in infected rabbits in prior experiments^10,11^. Samples were stored at -80°C until postmortem analysis unless otherwise specified.

#### Brain tissue processing and EV isolation

Initial optimization of EV isolation techniques was first performed with adult rabbit tissue for comparison of EV yield from freshly-collected and flash-frozen brain tissue. Adult rabbits had one hemisphere (∼4.5g/hemisphere) of brain tissue processed immediately after sample collection (i.e., freshly-collected) and the other hemisphere was flash-frozen and processed at a later time-point (i.e., flash-frozen) to allow for direct comparison from the same rabbit. To investigate the feasibility of extracting EVs from smaller brain samples (∼1g/hemisphere), one brain hemisphere was utilized from a 1-week-old healthy rabbit and from a infected rabbit 1-day post-infection. Samples were weighed and EV yield was corrected for tissue weight.

For all EV isolation, brains were minced and incubated in 10 mL of EV-release media^24^, composed of Neurobasal^TM^ medium (Gibco^TM^, cat# 21103049), supplemented with 1% GlutaMAX^TM^ supplement (Gibco^TM^, cat# 35050038) and 1% Antibiotic-Antimycotic (100X) (Gibco^TM^, cat# 15240062) at 37°C in a 5% CO2 environment for the 24h, except in the adult freshly-collected brain tissue that was incubated for 16h. The brain tissue supernatant was then clarified by differential centrifugation (300g for 10 min, 3000g for 10 min) and filtration through a 0.22 µm syringe filter (Millipore, cat# SLGPR33RS) to remove *M. tuberculosis*, cellular debris, apoptotic bodies, and larger microvesicles. The resulting supernatant contained the released EVs and underwent further purification through additional centrifugation (500g for 10 min, 3000g for 20 min, and 12,000g for 20 min) at room temperature (RT). The cleared supernatant was concentrated by centrifugation at 4,000g for 10 min at RT using Amicon Ultra-15 centrifugal filters (100 kDa MWCO) prior to size exclusion chromatography (SEC) using an automatic fraction collector (Izon Science, cat# AFC-V2). The concentrated supernatant was fractionated into an equal volume of 0.5 mL per fraction using a Gen 2 qEVoriginal/35 nm SEC columns (Izon Science, cat# ICO-35). Fractions were collected, concentrated to 150-200 µL using Amicon Ultra-0.5 mL centrifugal filters (10 kDa MWCO; Millipore Sigma, cat# UFC501008) and stored at -20°C until analysis. **Figure 1** illustrates the EV isolation protocol from brain tissue.

**Fig 1.**
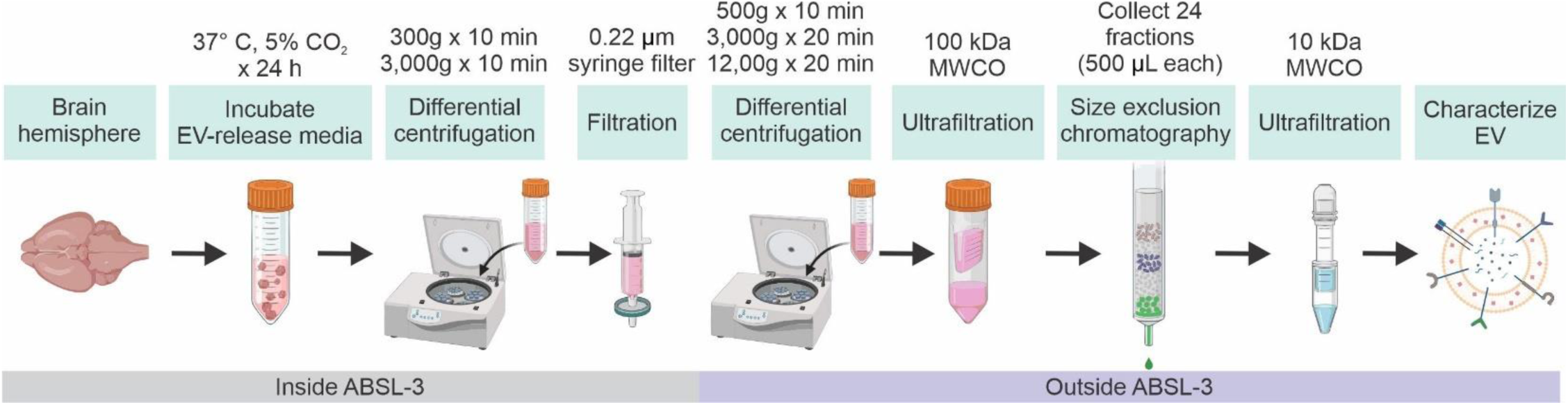
Workflow for the isolation of EVs from brain tissue. EV isolation steps for rabbit brain tissue performed inside and outside the animal biosafety-level 3 (ABSL-3) facility. Schematic was created in BioRender and CorelDRAW 2022.

### EV quantification by nanoflow cytometry (NanoFCM)

A NanoFCM Flow NanoAnalyzer (NanoFCM Co., Ltd) was used to measure EV concentration and size. Briefly, lasers were aligned and calibrated separately: fluorescent 250 nm silica nanoparticles at a concentration of 2.19 x 10^10^ (NanoFCM, cat# QS2503) were used for particle concentration, and a premixed silica nanosphere cocktail containing monodisperse nanoparticle populations of 68 nm, 91 nm, 113 nm, and 155 nm in diameter was used for size calibration (NanoFCM, cat# 516 M-Exo). PBS was used as the blank for background correction, and EVs were diluted 100-fold in PBS for particle quantification. Particle signal acquisition was performed for 1 min at a constant pressure of 1 kPa, with an event rate between 2,000 and 12,000 events/min. The side-scattering of each sample were analyzed using the NanoFCM Professional Suite V2.0 software.

### Transmission electron microscopy (TEM)

A subset of the EV samples was further characterized using TEM to visualize the particles and confirm expected EV morphology in EV-rich fractions. Samples (8 µL) were adsorbed to glow discharged (Electron Microscopy Sciences [EMS] GloQube) ultra-thin (UL) carbon coated 400 mesh copper grids (EMS, cat# CF400-Cu-UL), by floatation for 2 min. Grids were rinsed in 3 drops (1 min each) of Tris-buffered saline (TBS) buffer, negatively stained in 2 consecutive drops of 1% uranyl acetate (UA, aq.), and quickly aspirated. Grids were imaged on a Hitachi 7600 TEM operating at 80 kV with an AMT XR80 CCD (8 megapixels).

### Bicinchoninic acid (BCA) assay

The protein concentration of brain tissue lysates and EV samples was determined using a standard BCA Protein Assay (Thermo Fisher Scientific, cat# 23225) or Pierce Dilution-Free Rapid Gold BCA Protein Assay Kit (Thermo Fisher Scientific, cat# A55860) as indicated in the manufacturer’s manual. Briefly, samples were lysed using a ratio 1:1 of RIPA buffer (Thermo Fisher Scientific, cat# 89900,) supplemented with protease inhibitors (Thermo Fisher Scientific, cat# 78425,) on ice for 15 mins^25^. Ten microliters of diluted or undiluted samples were added to a 96-well plate, followed by 200 µL of the BCA/copper complex solution from the kit. Absorbance was measured at 564 nm for the standard BCA and 480 nm for the Pierce Dilution-Free Rapid Gold BCA in a microplate reader (Tecan Spark Multimode Microplate Reader) and protein concentration was determined against a bovine serum albumin (BSA) standard curve.

### SDS-PAGE and Western blots analysis

EV samples were lysed 1:1 with ice-cold RIPA lysis buffer (Thermo Scientific, cat # 89900) containing 1X Halt protease inhibitor cocktail (Thermo Scientific, cat# 78430) for 15 min. To denature and reduce the protein content in the samples, 1:4 ratio of Laemmli buffer (Bio-Rad, cat# 1610747) with dithiothreitol (DTT) was prepared and combined with the lysed EV samples at 95°C for 5 min. BCA protein assay was used to quantify protein. The denatured protein (10-20 µg) was loaded onto pre-cast polyacrylamide gels, 4-20% Mini-PROTEAN® TGX Stain-Free™ Protein Gels (Bio-Rad, cat# 4568093), with 1x Tris/Glycine/SDS (Bio-Rad, cat# 1610732) and subsequently transferred to nitrocellulose membranes using the Trans-Blot Turbo Mini 0.2 µm Nitrocellulose Transfer Packs (Bio-Rad, cat# 1704150). The membranes were blocked using a blocking buffer (SuperBlock T20 TBS Blocking Buffer, Thermo Scientific, cat# 37536) for 30 min at RT. The membranes were then incubated overnight at 4°C with specific primary antibodies (**Table 1**) diluted in blocking buffer. The next day, membranes were washed three times in TBS containing 0.1% Tween-20 (TBS-T) for 10 min and then incubated with appropriate secondary antibodies (**Table 2**) at RT for 1h. After the secondary antibody was removed with three washes of TBS-T, the membranes were incubated for 5 min with the following chemiluminescent substrates to visualize protein bands on the membranes; SuperSignal West Pico Maximum Sensitivity Substrate (Thermo Scientific, cat# 34580) or SuperSignal West Femto Maximum Sensitivity Substrate (Thermo Scientific, cat# 34095). The membranes were imaged with the ChemiDoc MP Touch imaging system (Bio-Rad) and analyzed using ImageJ (NIH) software.

**Table 1.** List of primary antibodies.

|  | **Primary antibody** | **Company** | **Catalog number** | **Western blot dilution** | **Dot blot dilution** |
| --- | --- | --- | --- | --- | --- |
| **EV-positive markers** | ALIX | Thermo Fisher Scientific | MA3-32773 | [1:1000] | - |
|  | Flotillin-1 | Thermo Fisher Scientific | MA5-35364 | [1:1000] | [1:500] |
|  | ATP1A3 | Thermo Fisher Scientific | MA3-915 | [1:1000] | [1:500] |
|  | GAPDH | Thermo Fisher Scientific | MA5-15738-HRP | [1:5000] | [1:500] |
| **EV-negative marker** | HSP70 | Abcam | Ab5439 | [1:1000] | [1:500] |
| **Cell-specific markers** | GFAP | Abcam | Ab4674 | [1:2000] | [1:500] |
|  | TMEM119 | Thermo Fisher Scientific | Cat# 66948-1 | [1:500] | [1:500] |
|  | GCP-II | GeneTex | GTX80151 | [1:1000] | [1:500] |
|  | NCAM1 | Biolegend | Cat# 304602 | [1:500] | [1:500] |
|  | CD11b | Biolegend | Cat. 101212 | - | [1:500] |
|  | GLS | Thermo Fisher Scientific | 6H5HK15 | [1:500] | [1:500] |

**Table 2.**
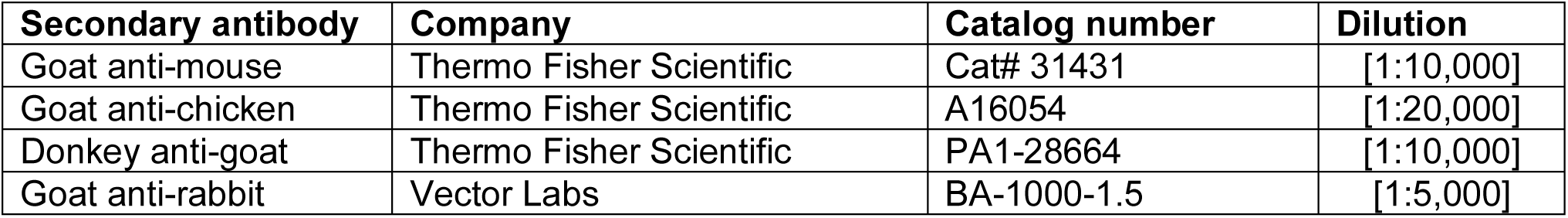
List of secondary antibodies.

| **Secondary antibody** | **Company** | **Catalog number** | **Dilution** |
| --- | --- | --- | --- |
| Goat anti-mouse | Thermo Fisher Scientific | Cat# 31431 | [1:10,000] |
| Goat anti-chicken | Thermo Fisher Scientific | A16054 | [1:20,000] |
| Donkey anti-goat | Thermo Fisher Scientific | PA1-28664 | [1:10,000] |
| Goat anti-rabbit | Vector Labs | BA-1000-1.5 | [1:5,000] |

### Dot-blot

Two microliters of EV lysate containing 100 ng of total protein were spotted onto nitrocellulose membranes. Separate membranes were prepared for each antibody tested. After air drying for 15 min, the membranes were blocked in blocking buffer for 30 mins at RT on a rocking platform. Primary antibodies (**Table 1**) were diluted in blocking buffer and incubated overnight at 4°C with individual membranes. The membranes were then washed three times with TBS-T, 10 min each wash, and then incubated with secondary antibody (**Table 2**) for 1h at RT. After washing, the signals were developed, imaged and analyzed as Western blots.

### Proteinase K protection assay

Proteinase K (PK) protection assay was utilized to identify which EV markers were on the surface or inside EVs. EV samples from three biological replicates per group (uninfected and *M. tuberculosis*-infected) were normalized to a concentration of 100 ng/µL total protein and divided into three treatment groups: (1) untreated intact EVs (vehicle control), (2) EVs treated with PK (Thermo Fisher Scientific, cat# EO0491) alone, and (3) EVs pre-treated with Triton X-100 (Sigma-Aldrich, cat# T8787-50ML) followed by PK. For permeabilization, EVs were incubated with 1% (v/v) Triton X-100 on ice for 15 min. Samples were then treated with PK (0.5 µg/mL final concentration) for 1h at 37°C. PK was inactivated by the addition of phenylmethylsulfonyl fluoride (PMSF; Cell Signaling Technology, cat# 8553S) to a final concentration of 2 mM, followed by incubation at RT for 15 min. All reactions were adjusted to a final volume of 40 µL using PBS. A 1 µL aliquot from each sample (containing 100 ng EV protein) was then applied to a nitrocellulose membrane for dot blot analysis to determine if EV-associated proteins were digested by PK alone (i.e., surface-associated proteins) or required permeabilization with Triton X-100 (i.e., internalized proteins).

### Comparative proteomics of brain-derived EVs

After EV isolation, a subset of EVs (n = 5/group) was sent for comparative proteomic analysis. EVs were solubilized in 1% SDS and sonicated. Twenty micrograms of total protein per sample were digested with trypsin/lysC, desalted with a C18 column, and reconstituted in 0.1% formic acid. LC-MS/MS analysis was conducted using a Thermo Orbitrap Exploris 240 Mass Spectrometer and a Thermo Vanquish Neo RSLCnano system (Poochon Scientific). Peptide mixture from each sample was loaded onto a peptide trap cartridge at a flow rate of 5 µL/min. The trapped peptides were eluted onto a reversed-phase Easy-Spray column using a linear gradient of acetonitrile (3-36%) in 0.1% formic acid. The elution duration was 110 min at a flow rate of 0.3 µL/min. Eluted peptides from the EasySpray column were ionized and sprayed into the mass spectrometer, using a Nano-EasySpray Ion Source (Thermo) under the following settings: spray voltage, 1.6 kV, capillary temperature, 275°C. Raw data files acquired from each sample were searched against *Oryctolagus cuniculus* and *M. tuberculosis* (strain ATCC 25618 / H37Rv) protein sequences databases using the Proteome Discoverer 2.5 software (Thermo, San Jose, CA) based on the SEQUEST algorithm. Carbamidomethylation (+57.021 Da) of cysteines was fixed modification, and Oxidation Met and Deamidation Q/N-deamidated (+0.98402 Da) were set as dynamic modifications. The minimum peptide length was specified to be five amino acids. The precursor mass tolerance was set to 15 ppm, whereas fragment mass tolerance was set to 0.05 Da. The maximum false peptide discovery rate was specified as 0.01. The resulting Proteome Discoverer Report contains all assembled proteins with peptides sequences and peptide spectrum match counts (PSM#) and MS1 peak area abundance. Protein quantification/normalization used the normalized spectral abundance factors (NSAFs) method to calculate the protein relative abundance^26,27^.

### Bioinformatic analysis

Enrichment of subcellular localization terms was performed using the Database for Annotation, Visualization, and Integrated Discovery (DAVID; v2023q4, https://david.ncifcrf.gov/), with the *Homo sapiens* proteome used as background. Analysis was limited to Gene Ontology Cellular Component (GO:CC) terms. Statistical significance was determined using Fisher’s exact test with Benjamini-Hochberg correction (adjusted *P* < 0.05). The top 10 most significantly enriched cellular components for each group were visualized as bar graphs using Prism GraphPad.

Pathway and cell-type enrichment analyses were conducted using Enrichr (https://maayanlab.cloud/Enrichr/). CellMarker 2024 databases were queried for all EV protein groups to validate cell-type associations. For pathway analysis, Reactome Pathways 2024 was used for upregulated and downregulated proteins, while WikiPathways 2024 was applied specifically to proteins exclusively detected in EVs from infected rabbits. Enrichment results were ranked by adjusted p-value, and the top terms were visualized in Prism GraphPad.

Protein–protein interaction networks were generated using the STRING database (v11.5; https://string-db.org/) with a minimum interaction confidence score of 0.4. Networks were visualized and clustered in Cytoscape (v3.9.1) using the clusterMaker plugin. Functional enrichment analysis was performed on each module using STRING’s built-in enrichment tools. Enriched biological processes and pathways were categorized and sorted based on their corresponding cluster membership.

To evaluate the overlap of identified brain EV proteins with known EV cargo, we compared all upregulated, downregulated, and proteins exclusively detected in *M. tuberculosis*-infected samples (i.e., infection-specific) to the Vesiclepedia database using FunRich (http://www.funrich.org). Venn diagrams were generated to visualize shared and unique protein subsets between our dataset and established EV protein signatures.

### Statistical analysis

All data were analyzed using Prism 10 software (GraphPad). Data are expressed as mean ± SD. Comparisons of parametric data were performed using t-tests for single comparisons and ordinary one-way analysis of variance (ANOVA) followed by Tukey’s multiple comparison tests for multiple comparisons. *P* values ≤ 0.05 were considered statistically significant. Empirical Bayesian-modified t-tests were performed with the linear modeling of microarrays (LIMMA) R package. Proteins were considered significantly differentially expressed at p < 0.05 and/or q < 0.05 (Benjamini-Hochberg correction). Principal component analysis (PCA) was performed on proteins with p < 0.05. Heatmaps were generated using ComplexHeatmap, with row clustering based on Euclidean distance. Volcano plots highlight proteins passing the significance threshold and labeled based on top 50 p-values and top 20 absolute log₂ fold changes. All visualizations were created in R using ggplot2, ggrepel, and ComplexHeatmap packages. Functional relationships between proteins were collated with stringApp 2.0.1 embedded in Cytoscape 3.10.1, using a string score of 0.7 as the evidence threshold for association. Figures and graphs were created with BioRender and CorelDRAW 2022.

## RESULTS

### Adaptation of the EV release method for EV isolation from fresh and flash-frozen rabbit brain tissue

To our knowledge, EV isolation from rabbit brain tissue has not been previously reported. To address this gap, we tested whether the “spontaneous EV release” method, originally developed for mouse and human brain tissue^24^, could be adapted for use in rabbits. This method was combined with standard EV purification steps including differential centrifugation, ultrafiltration, and size-exclusion chromatography (SEC), as illustrated in **Figure 1**.

Initial feasibility testing was performed using brain tissue from uninfected adult rabbits. Given the constraints of collecting rabbit brain tissue under biosafety level-3 (BSL-3) conditions, immediate postmortem incubation in EV release media would limit samples obtained. Therefore, we evaluated whether the protocol could be adapted for flash-frozen tissue without compromising EV integrity. A single adult rabbit brain was bisected; one hemisphere was processed fresh, while the other was flash-frozen for one week prior to EV isolation (**S1 in supplementary figure files**). Following SEC, 13 fractions (F) were collected and analyzed by BCA protein assay and NanoFCM. In fresh tissue, EVs were enriched in fractions F1–F3, while in frozen tissue, enrichment extended through F1–F4. Visual inspection by TEM revealed abundant EV-like particles in fractions F1–F3 of fresh samples and F1–F5 of frozen samples. TEM also revealed the expected cup-shaped morphology, with diameters ranging from 50–200 nm. Mean particle size was comparable between conditions, and EV yield, normalized to tissue weight, was slightly higher in frozen samples. Altogether, these findings support the use of frozen rabbit brain tissue for EV isolation.

### Application of the EV isolation protocol to neonatal rabbits following M. tuberculosis infection

Building on our results in adult rabbits, we next applied the optimized “EV release” method to brain tissue collected from one-week-old rabbits to test the EV yield from smaller samples. EVs were isolated from an uninfected rabbit and an infected littermate sacrificed 24 hours post-infection (**S2 in supplementary figure files**). Similar to findings in adults, EVs were enriched in fractions F1–F5 in both uninfected and infected samples. This was supported by NanoFCM analysis of all 13 fractions and visual inspection of pooled EV-rich fractions by TEM, which confirmed the presence of vesicles with characteristic morphology and size (**S2A-D**).

To further characterize EVs from one-week-old rabbits, we performed Western blot analysis on SEC fractions for the EV markers Hsp70 and Alix, along with GCPII, a protein associated with microglia and astrocytes. These proteins were enriched in fractions F1–F5, consistent with NanoFCM and TEM data (**S2E,F**). We then pooled fractions F1–F5 from each sample to compare EV cargo between the uninfected and *M. tuberculosis*-infected groups. Western blotting revealed similar GCPII levels in both groups, while Hsp70 and ATP1A3 appeared more abundant in EVs from the infected rabbit (**S2G**). Given the limited sample size (n=1 per group), these trends should be interpreted cautiously. Having validated that fractions F1–F5 reliably contain EVs in both adult and neonatal rabbit brains, we adopted this approach for EV isolation in our subsequent 21-day post-infection study.

### M. tuberculosis infection increases EV release in brain tissue of young rabbits

To determine whether *M. tuberculosis* infection influences brain EV biogenesis during disease progression, we analyzed brain EVs isolated from rabbits 21 days after infection. TEM revealed typical cup-shaped vesicles in both uninfected and infected rabbits (**Figure 2A**), and NanoFCM analysis showed similar size distributions and mean particle diameters across groups (**Figure 2B,C**). Despite similar morphology, the total EV concentration per 100 mg of brain tissue was significantly higher in *M. tuberculosis*-infected rabbits (**Figure 2D**). Despite the significant increase in EV concentration in infected brain tissue, total EV protein levels were comparable to uninfected (**Figure 2E**), suggesting elevated vesicle secretion without a corresponding increase in protein content. These findings suggest that *M. tuberculosis* infection enhances EV release in the brain, potentially reflecting altered vesicle secretion dynamics in response to infection.

**Fig 2.**
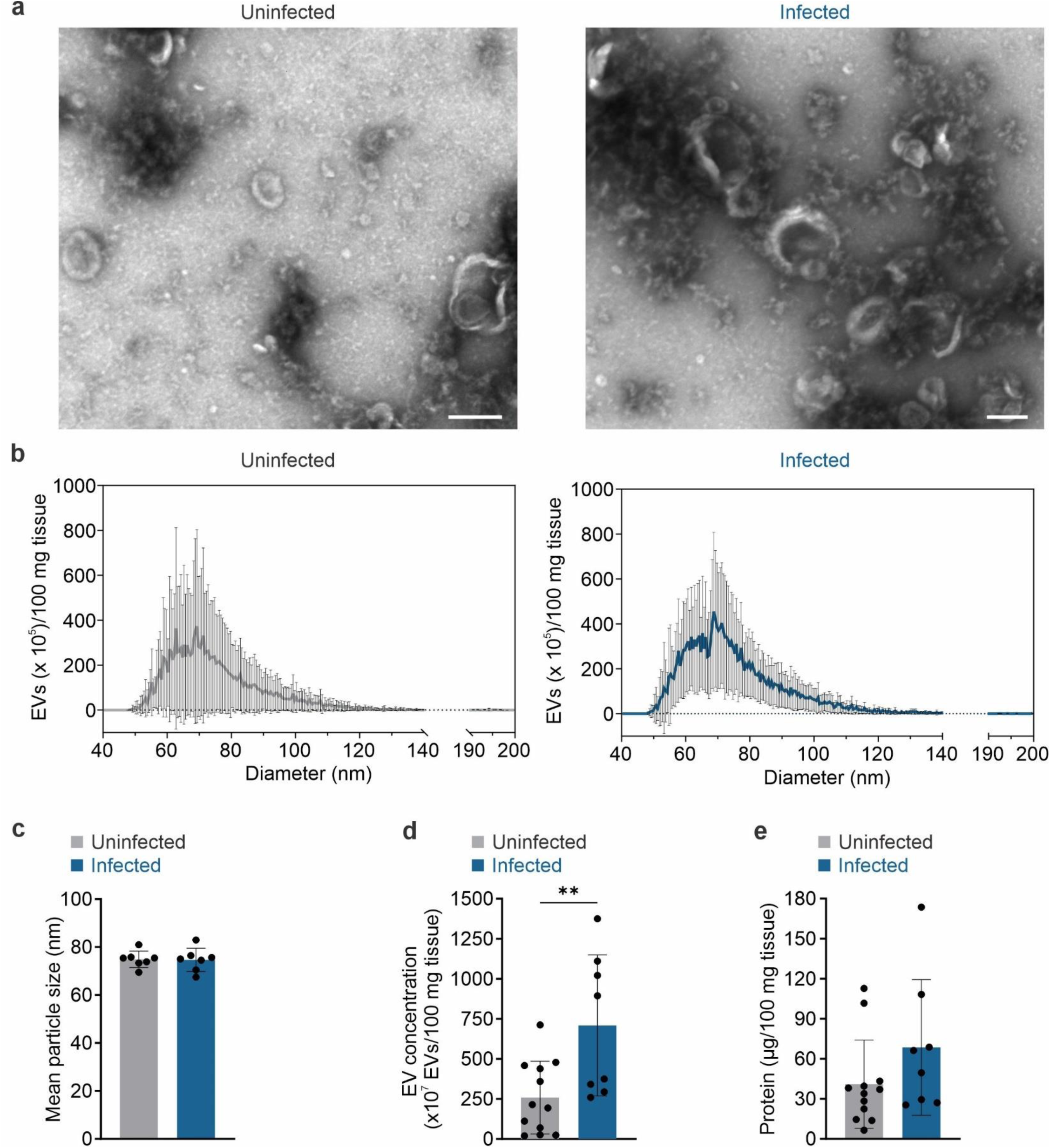
Brain-derived EVs from young rabbits with TB meningitis and age-matched healthy controls. EVs were isolated from brain tissue from young rabbits with TB meningitis and age-matched healthy controls 3-weeks after injection with *M. tuberculosis* (*M. tuberculosis*-infected) or PBS (uninfected) respectively. **a** Negative-stain transmission electron microscopy (TEM) visualized brain-derived EVs from uninfected (left) and *M. tuberculosis*-infected (right) rabbits. Scale bar = 200 nm. **b** Size distribution determined by Nanoflow cytometry (NanoFCM) for uninfected (left) and *M. tuberculosis*-infected (right) brain-derived EVs. EV quantification was normalized per 100 mg of brain tissue. n = 7 animals/group. **c** Mean particle size of brain-derived EVs. **d** EV concentration measured by NanoFCM and normalized per 100 mg of brain tissue. **e** Protein concentration measured by BCA of brain-derived EVs. Data are represented as mean ± standard deviation (SD). Statistical comparisons were made using unpaired two-tailed t-test. \*\**P* < 0.01

### Distinct protein signatures define EVs from M. tuberculosis-infected brains

To investigate the impact of *M. tuberculosis* infection on brain-derived EVs, we performed label-free quantitative proteomic analysis of EVs isolated from the brain tissue of a subset of infected and uninfected rabbits. Principal component analysis (PCA) revealed clear group separation, indicating distinct proteomic signatures between infected and uninfected brain-derived EVs (**Figure 3A**). Differential expression analysis identified a panel of proteins significantly enriched or downregulated in brain-derived EVs during *M. tuberculosis* infection. A volcano plot (**Figure 3B**) visualized proteins that were significantly upregulated or downregulated in brain-derived EVs from infected compared to uninfected rabbits (p < 0.05, |log₂FC| > 1). Hierarchical clustering in the form of a heatmap of the top 30 proteins that passed a false discovery rate (FDR) threshold of q < 0.05 revealed clear group-specific clustering, with several host response proteins elevated in the *M. tuberculosis* infection (**Figure 3C**). Additionally, a number of proteins were significantly upregulated (n = 321), downregulated (n = 248), or uniquely expressed (i.e., infection-specific, n = 184) in brain-derived EVs from *M. tuberculosis*-infected rabbits, with a large proportion of the differentially expressed and exclusive proteins overlapped with known EV proteins in the Vesiclepedia database, further validating the vesicular origin of the dataset (**Figure 3D**).

**Figure 3.**
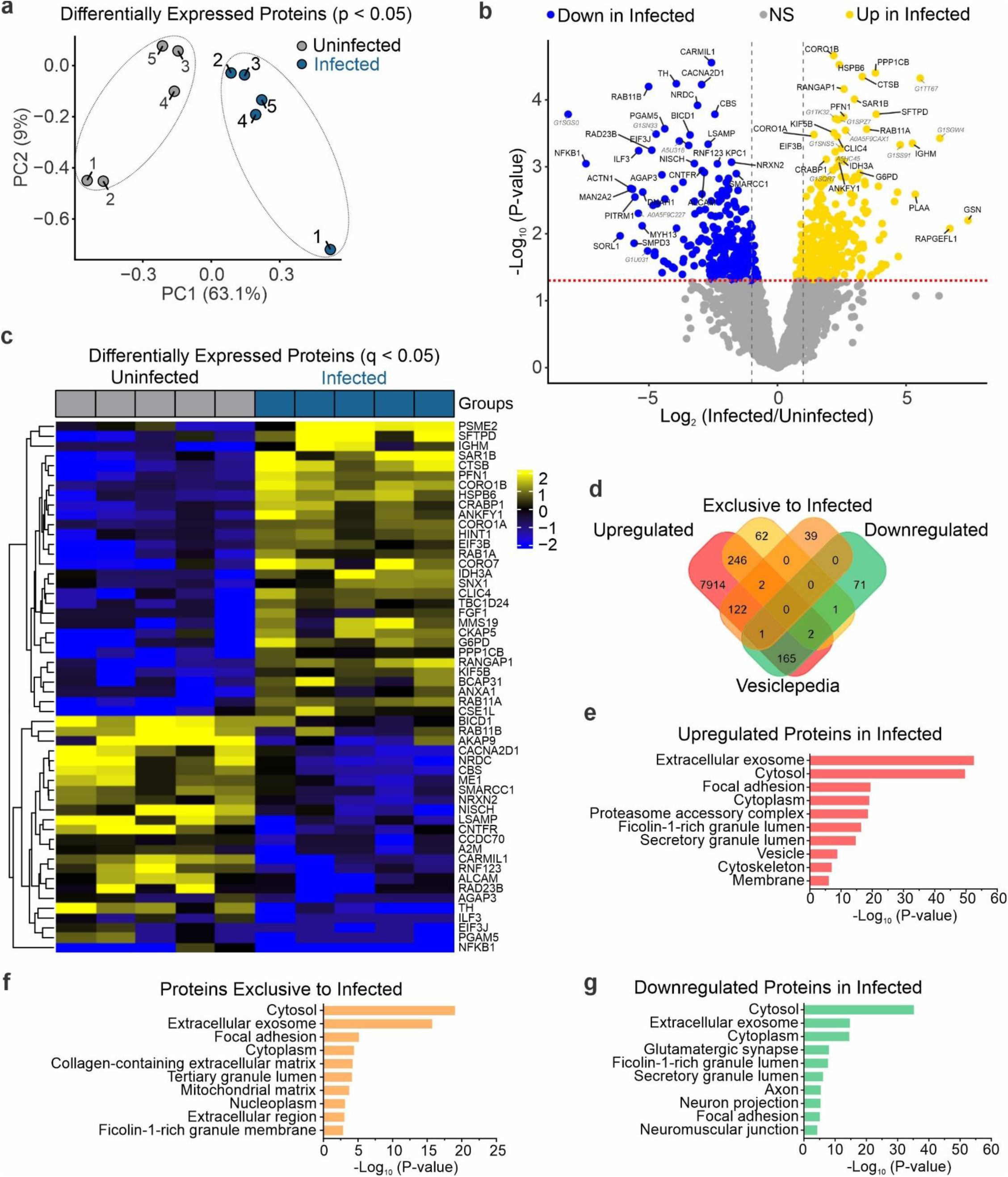
Protein profiling of brain-derived EVs. **a** Principal Component Analysis (PCA) of significantly differentially expressed proteins (p < 0.05), showing clear separation between uninfected control and *M. tuberculosis*-infected EV samples. **b** Volcano plot showing differentially expressed EV proteins. Yellow and blue points denote proteins significantly upregulated or downregulated in *M. tuberculosis*-infected EVs, respectively (p < 0.05 and log₂FC > 1). Select top proteins by p-value and log₂ fold change are labeled. **c** Heatmap of all proteins with FDR-adjusted q-values < 0.05. **d** Venn diagram illustrates common proteins that are upregulated, downregulated (*M. tuberculosis*-infected/uninfected), or exclusively detected in *M. tuberculosis*-infected brain tissue, compared to the Vesiclepedia database. The analysis was performed using FunRich. **e-g** Database for Annotation, Visualization and Integrated Discovery (DAVID) was used to determine the most enriched cellular compartments associated with proteins significantly enriched or exclusively present in brain-derived EVs from infected rabbits. Ten of the most enriched terms (based on p-value) in each category are displayed.

To evaluate the subcellular origin of EV cargo alterations, we performed cellular compartment enrichment analysis (DAVID) on upregulated, downregulated, and infection-specific proteins in brain-derived EVs from infected rabbits (**Figure 3E–G**). The top ten enriched categories across all groups included extracellular exosome, cytoplasm, and cytosol, consistent with canonical EV cargo profiles. Upregulated proteins showed additional enrichment in components related to protein turnover and trafficking, including proteasome accessory complexes, focal adhesion sites, and secretory granule lumens. In contrast, downregulated proteins were enriched in neuron-associated compartments, such as the glutamatergic synapse, axon, and neuromuscular junction, suggesting lower inclusion of neurostructural or signaling proteins. Proteins exclusively detected in brain-derived EVs from infected rabbits were linked to a broader range of compartments, including the extracellular matrix, mitochondrial matrix, and nucleoplasm, reflecting infection-driven packaging of EV proteins from multiple intracellular sources.

### Host immune and stress-response proteins are enriched in brain-derived EVs during M. tuberculosis infection

To investigate the biological relevance of proteins upregulated in brain-derived EVs from infected rabbits, we performed *in silico* enrichment and protein interaction network analyses. Enrichr CellMarker 2024 analysis revealed strong enrichment for markers of microglia, astrocytes, and immune cell populations (**Figure 4A**), suggesting a shift in the cellular origin or selective loading of EV cargo during infection. Reactome Pathways 2024 analysis of the same upregulated protein set identified enrichment in immune and vesicle-related processes, including neutrophil degranulation, antigen presentation, and endosomal transport (**Figure 4B**). STRING protein–protein interaction analysis clustered the upregulated proteins into five distinct functional groups (**Figure 4C**). Cluster-specific enrichment analysis (**Figure 4D**) revealed biological processes associated with chaperone-mediated protein folding (Cluster 1), RNA processing (Cluster 2), cytoskeletal dynamics (Cluster 3), vesicle trafficking (Cluster 4), and proteasome-associated protein degradation (Cluster 5). Together, these results indicate that brain-derived EVs from infected brain tissue are enriched in host proteins associated with immune activation, stress responses, and intracellular transport, potentially reflecting adaptive host responses to infection.

**Figure 4.**
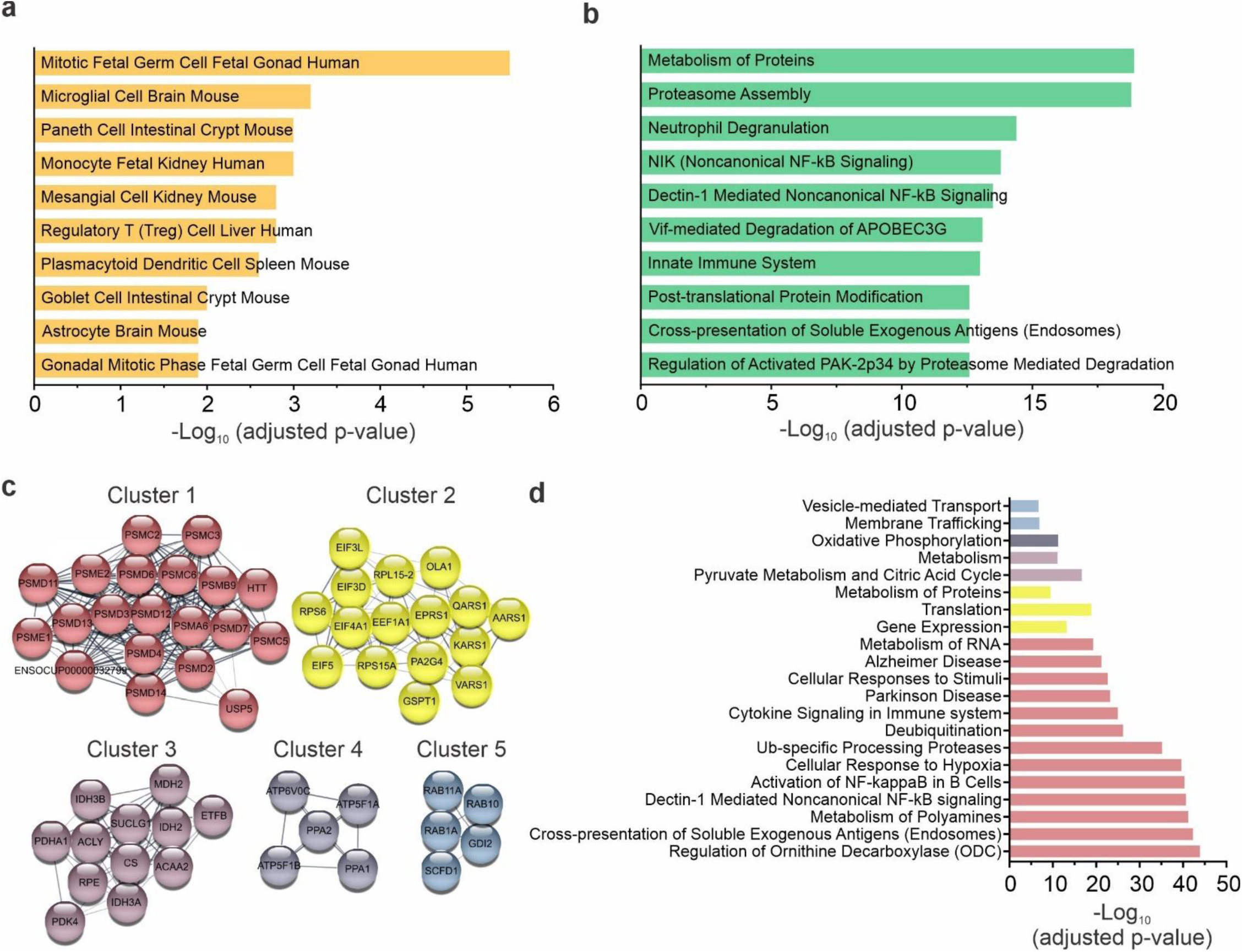
Cell type and biological pathways associated with proteins upregulated in EVs from infected brain tissue. **a-b**. Top 10 significant (**a**) Enrichr GO cellular component 2024 and (**b**) Enrichr Reactome Pathways 2024 of proteins upregulated in *M. tuberculosis*-infected EV samples, sorted by p-value. **c** STRING functional protein association network clusters. Nodes represent proteins, and edges indicate predicted functional interactions based on known associations. **d** STRING functional enrichment pathways are categorized and sorted according to the corresponding cluster displayed in (**c**), highlighting the biological processes and pathways enriched within each cluster.

Several enriched proteins during infection have well-documented roles in host responses against *M. tuberculosis* and in CNS pathology. Coronin-1A (CORO1A), a cytoskeletal regulator that prevents phagosome maturation to enable *M. tuberculosis* survival^28,29^, is also elevated in pulmonary and extrapulmonary TB, supporting its potential as a clinical biomarker^30^. Similarly, cathepsin B (CTSB), a lysosomal protease that promotes inflammasome activation and IL-1β release, is also elevated in the lungs of *M. tuberculosis*-infected mice and rabbits and in the plasma of patients with active TB^31^, underscoring its systemic role in infection. Surfactant protein D (SFTPD), which binds to mycobacterial glycolipids^32^ and facilitates bacterial clearance^33^ was also elevated in EVs from infected brain. Though traditionally considered lung-specific, SFTPD has recently been detected in human CSF and CNS tissue under both healthy and pathological conditions^34,35^, suggesting a role in CNS immune surveillance. Annexin A1 (ANXA1), an anti-inflammatory mediator that promotes resolution of inflammation and clearance of apoptotic cells^36^, was also elevated during *M. tuberculosis* infection. In the CNS, annexin A1 is expressed by microglia and has been shown to reduce neuroinflammation^37^ and ameliorate amyloid beta pathology in Alzheimer’s disease models^38^, while its absence (i.e., annexin A1-deficient mice) leads to higher *M. tuberculosis* burden and disorganized granulomas^39^. Its presence in brain-derived EVs likely reflects a host-protective response to limit tissue damage in TB meningitis. Lastly, brain-derived EVs were also enriched with the immunoglobulin M (IgM) heavy chain (IGHM), consistent with early activation of humoral immunity during infection. Notably, IgM is elevated in CSF in TB meningitis patients and correlated with disease severity, poor prognosis and treatment response^40^, suggesting its potential as a dynamic biomarker for disease monitoring.

### Infection-specific brain-derived EVs reflect microglial activation and interferon-driven immune signaling

To characterize infection-specific protein cargo selectively packaged into EVs, we analyzed proteins exclusively detected in brain-derived EVs from infected rabbits. Enrichr CellMarker 2024 analysis revealed strong enrichment for microglial, monocyte, neutrophil, and myeloid-derived suppressor cell markers (**Figure 5A**), suggesting increased EV contributions from innate immune cells in the infected brain. WikiPathways 2024 analysis further revealed enrichment in immune and infection-related pathways, including the immune response to TB, type I interferon (IFN) signaling, microglial phagocytosis, and viral infection–related signaling networks (**Figure 5B**). STRING interaction network analysis grouped these exclusive proteins into five distinct clusters (**Figure 5C**). STRING enrichment analysis of each cluster (**Figure 5D**) highlighted pathways such as IFN-stimulated gene expression (Cluster 1), nucleotide metabolism (Cluster 2), DNA damage response and base excision repair (Cluster 3), protein folding and cytokine responses (Cluster 4), and IL-17 signaling (Cluster 5). These data predict that EVs in the infected brain selectively incorporate proteins associated with innate immunity, IFN interferon signaling, and host defense, reflecting a potential role for brain-derived EVs in amplifying immune-activating signals during *M. tuberculosis* infection.

**Figure 5.**
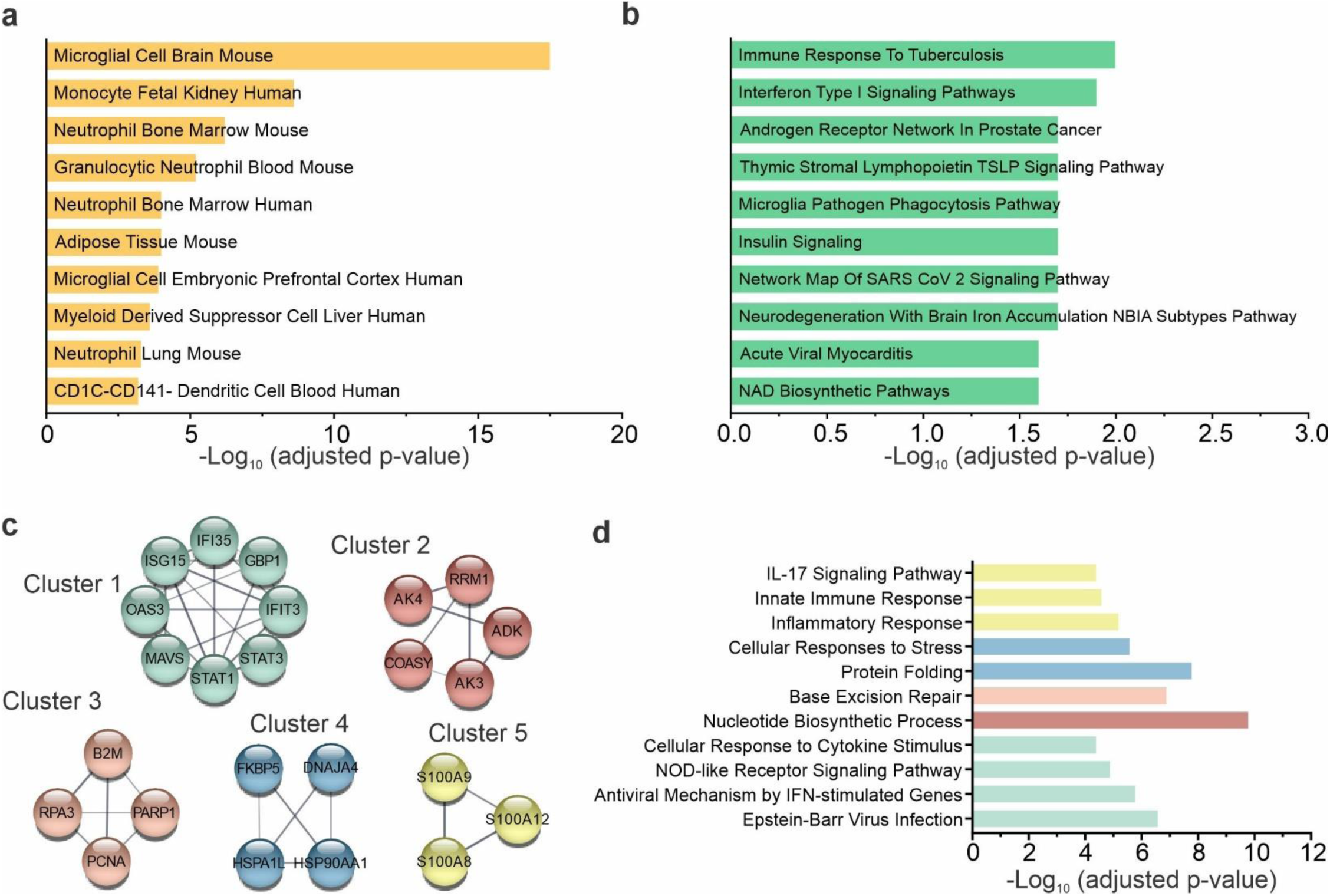
Cell type and biological pathways associated with proteins detected exclusively in EVs from infected brain tissue. **a-b**. Top 10 significant (**a**) Enrichr GO cellular component 2024 and (**b**) Enrichr WikiPathways 2024 of proteins exclusive to *M. tuberculosis*-infected EV samples (i.e. infection-specific), sorted by p-value. **c** STRING functional protein association network clusters. Nodes represent proteins, and edges indicate predicted functional interactions based on known associations. **d** STRING functional enrichment pathways are categorized and sorted according to the corresponding cluster displayed in (**c**), highlighting the biological processes and pathways enriched within each cluster.

We also identified several inflammatory proteins uniquely expressed in EVs from infected brains. Profilin-1 (PFN1), an actin-binding protein implicated in microglial activation, vesicular trafficking, and pro-inflammatory responses after CNS injury^41–43^, was only detected during *M. tuberculosis* infection. Glycoprotein nonmetastatic melanoma protein B (GPNMB), a marker of disease-associated microglial activation^44^, galectin-3 (LGALS3), a known regulator of microglial activity and neuroinflammation^44,45^ and, lactotransferrin (LTF), which is released by activated microglia and has immunomodulatory and antimicrobial activity, were also only present in EVs from infected rabbits. Together these proteins support the central role of microglial activation in TB meningitis. The interferon-stimulated proteins ISG15, OAS3, IFIT3, and STAT1 were also enriched, consistent with the type I interferon bias in active TB^46–48^. Within this group, GBP1 was notable for its IFN-inducibility, anti-mycobacterial function, and emerging value as a TB biomarker^49^. Additionally, we detected gasdermin D (GSDMD), a protein cleaved during NLRP3 inflammasome activation to trigger pyroptosis and IL-1β release^50^, suggesting a vesicle-associated mechanism for amplifying inflammasome signaling in the CNS. Finally, the neutrophil alarmins S100A8 and S100A9 were uniquely present, aligning with their elevated CSF levels in TB meningitis patients and their established role in neutrophil-driven inflammation^51^.

### M. tuberculosis infection reduces neuronal and neuroprotective protein cargo in brain-derived EVs

To gain insight into host processes downregulated during *M. tuberculosis* infection, we analyzed proteins significantly underrepresented in brain-derived EVs from infected rabbits. CellMarker 2024 enrichment revealed that these proteins were highly associated with neuronal and glial cell types, including human neurons, astrocytes, and oligodendrocytes (**Figure 6A**), suggesting reduced incorporation of proteins key for maintenance of brain structure, myelination and metabolic balance into brain-derived EVs during infection. Pathway analysis using Enrichr Reactome Pathways 2024 indicated that downregulated proteins were involved in key neurodevelopmental and intracellular signaling pathways, such as L1CAM interactions, axon guidance, and mitochondrial apoptosis signaling (**Figure 6B**). Protein–protein interaction analysis via STRING revealed six distinct functional clusters among downregulated proteins (**Figure 6C**). STRING enrichment analysis (**Figure 6D**) showed that Cluster 1 was enriched in metabolic and carbohydrate-related pathways, Cluster 2 in protein binding and apoptotic regulation, and Cluster 3 in amyloid regulation and signal transduction. Cluster 4 included vesicle biogenesis and trafficking components, Cluster 5 encompassed cholesterol biosynthesis and membrane-associated pathways, and Cluster 6 was strongly associated with nervous system development and axon guidance. Together, these findings suggest that *M. tuberculosis* infection leads to reduced EV packaging of neurostructural, synaptic, and intracellular trafficking proteins, potentially reflecting disruption of neuronal function and neurodevelopmental signaling in the infected brain.

**Figure 6.**
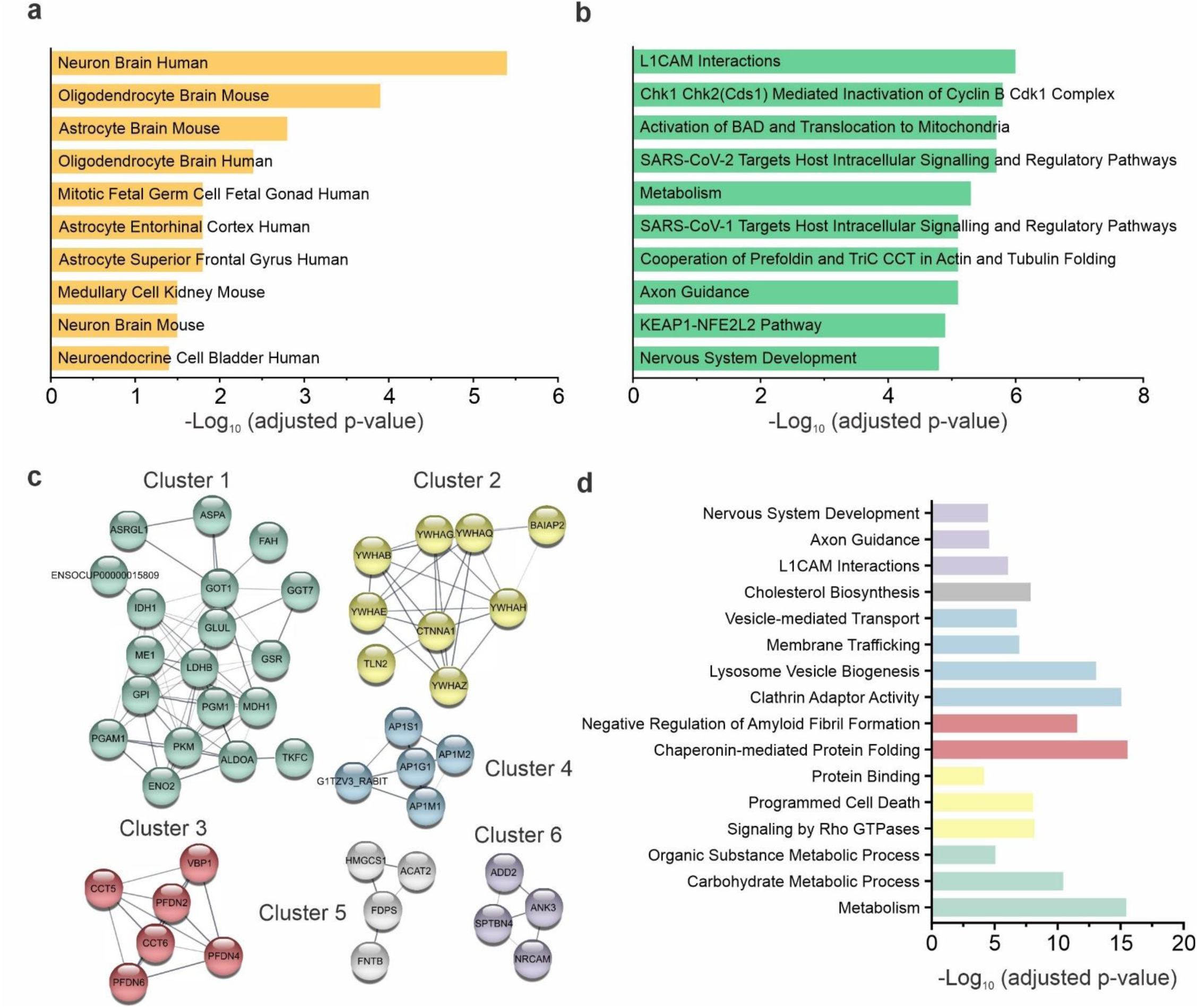
Cell type and biological pathways associated with proteins downregulated in EVs from infected brain tissue. **a-b**. Top 10 significant (**a**) Enrichr CellMaker component 2024 and (**b**) Enrichr Reactome Pathways 2024 of proteins downregulated in *M. tuberculosis*-infected EV samples, sorted by p-value. **c** STRING functional protein association network clusters. Nodes represent proteins, and edges indicate predicted functional interactions based on known associations. **d** STRING functional enrichment pathways are categorized and sorted according to the corresponding cluster displayed in (**c**), highlighting the biological processes and pathways enriched within each cluster.

While most infection-specific changes reflected immune activation, several proteins important for neuronal integrity and neuroprotection were significantly reduced in EVs from infected rabbits. SORL1, an endosomal sorting receptor implicated in amyloid precursor trafficking and Alzheimer’s disease, was decreased, consistent with endosomal dysfunction seen in neurodegeneration^52^. PITRM1, a mitochondrial protease responsible for degrading amyloid-β peptides, was diminished as well. Interestingly, reduced PITRM1 activity has been linked to amyloid accumulation and Alzheimer’s-like pathology in animal models^53^ and amyloid deposition has been seen in our rabbit model of TB meningitis. Tyrosine hydroxylase (TH), the rate-limiting enzyme in dopamine synthesis and a marker of dopaminergic neuron integrity, was also reduced, mirroring molecular patterns observed in Parkinson’s disease^54^. Together, these depletions suggest disruption of neuroprotective signaling and mitochondrial homeostasis during TB meningitis, offering a potential mechanism for the long-term neurological sequelae observed in survivors^3,4^.

### Proteomic trends in cell-specific EV protein expression

Intrigued by the CellMarker 2024 cell type enrichment data, we classified differentially expressed brain-derived EV proteins by their predicted cellular origin. This analysis revealed distinct patterns across neurons, astrocytes, and microglia (**Figure 7A**). While several proteins associated with microglia and astrocytes were elevated in brain-derived EVs from infected rabbits, a substantial number of neuron-associated proteins, including NCAM1, SYT1, and GRIA1, were significantly reduced. These trends suggest a shift in EV contributions or cargo selection away from neuronal sources during infection.

**Figure 7.**
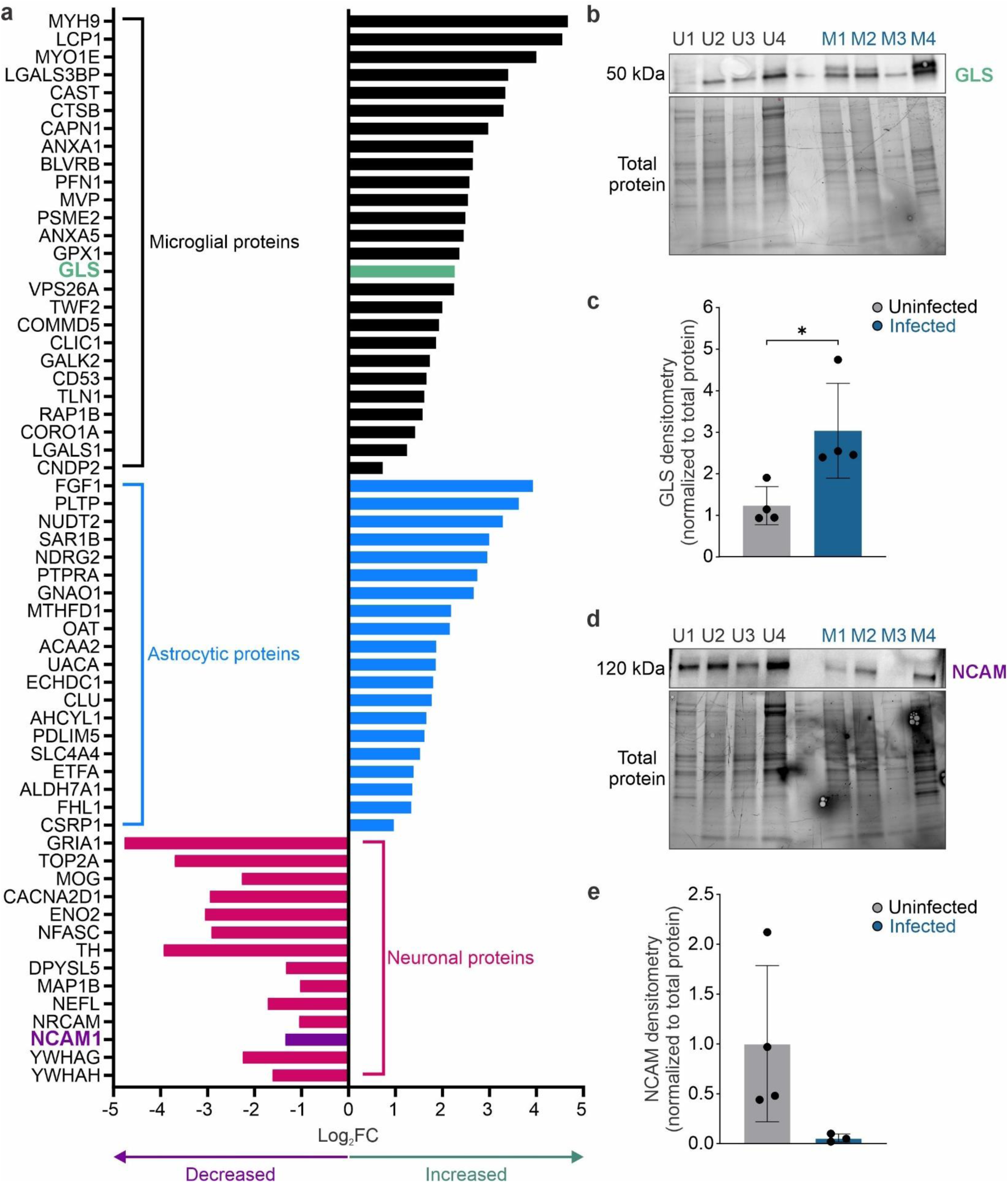
Differential expression of cell type–associated proteins in brain-derived EVs. **a** Log2 fold change (Log2FC) of brain-derived EV proteins classified by brain cell type based on curated annotations (Enrichr CellMarker 2024). Proteins associated with microglia (black), astrocytes (blue), and neurons (pink) are shown; positive values indicate increased enrichment in *M. tuberculosis*-infected EV samples. GLS (mitochondrial enzyme expressed on activated microglia) expression was increased and NCAM1 (neuronal adhesion molecule) expression was decreased in brain-derived EVs. **b,c** Increased GLS expression in brain-derived EVs from infected samples (M, blue) compared to uninfected control (U, black/grey) validated with (**b**) Western blot and (**c**) corresponding densitometry. GLS was significantly increased in infected EV samples (*P* < 0.05). n = 4 rabbits/group. **d,e** Decreased NCAM1 expression in brain-derived EVs from *M. tuberculosis*-infected rabbits validated with (**d**) Western blot and (**e**) corresponding densitometry, though not statistically significant. n = 4 rabbits/group. All protein levels were normalized to total protein loading. Data are presented as mean ± SD from biologically independent samples. Statistical comparisons were made using unpaired two-tailed t-test. \**P* < 0.05.

Next, we used Western blot analysis to validate the proteomics data of selected cell-type–specific proteins in brain-derived EVs from uninfected and infected rabbit brain tissue (**Figure 7B–E, S3 in supplementary figure files**). ATP1A3 (a neuronal ion transporter) and GFAP (an astrocytic intermediate filament protein) were detected at similar levels across both conditions, indicating no significant changes in their EV representation (**S3 in supplementary figure files**). Likewise, the microglial marker TMEM119 remained stable between groups, as did GCPII, a glia-associated protein, suggesting that these specific glial EV proteins represent a portion of EV cargo preserved at this stage of infection (**S3 in supplementary figure files**). Interestingly, expression of glutaminase (GLS), a mitochondrial enzyme predominantly expressed in neurons under homeostatic conditions and upregulated in activated microglia under pathological conditions^55^, was significantly increased in brain-derived EVs from infected brains (**Figure 7B,C**). This elevation may reflect enhanced glutaminolytic activity or metabolic reprogramming in glial cells during neuroinflammation, contributing to altered EV cargo profiles. In contrast, NCAM1, a neuronal adhesion molecule, showed a pronounced reduction in EVs from infected brains, in line with the proteomic data, though the change did not reach statistical significance (**Figure 7D,E**). These findings suggest that *M. tuberculosis* infection results in selective remodeling of EV protein cargo, characterized by reduced inclusion of some neurostructural proteins and increased representation of metabolically active or stress-related enzymes, potentially reflecting altered EV biogenesis, intercellular signaling, and metabolic adaptation within the infected CNS environment.

### Brain-derived EVs contain an array of internal and surface-associated protein markers

To investigate the sub-vesicular localization of specific proteins within the brain-derived EVs, we performed PK protection assays (**Figure 8A,B**). In this assay, surface-associated EV proteins are degraded by PK, while proteins encapsulated within the vesicle lumen remain protected unless the membrane was permeabilized by detergent (i.e., Triton X-100). Dot blot analysis of brain-derived EVs from uninfected rabbits revealed that the microglial markers CD11b and TMEM119, as well as the astrocyte marker GFAP and the metabolic enzyme GAPDH, were degraded following PK treatment even in the absence of detergent (**Figure 8C**). This suggests that these proteins are primarily associated on the EV surface, rendering them accessible to proteolytic cleavage. In contrast, ATP1A3, NCAM, GCPII, and GLS were resistant to protease treatment in the absence of detergent but were degraded when the EV membrane was permeabilized with Triton X-100 (**Figure 8D**). These findings indicate that these proteins are predominantly localized within the EV lumen or associated with the inner membrane. A similar protection pattern was observed in EVs isolated from infected rabbit brain tissue, suggesting that infection does not alter the internal localization of these particular proteins. Together, these data support the distinct sub-vesicular compartmentalization of select EV proteins and reveal that both uninfected and infected brain-derived EVs harbor a mixture of internal and surface-exposed protein markers. This information could inform the selection of surface-associated EV proteins for EV capture or targeting applications.

**Figure 8.**
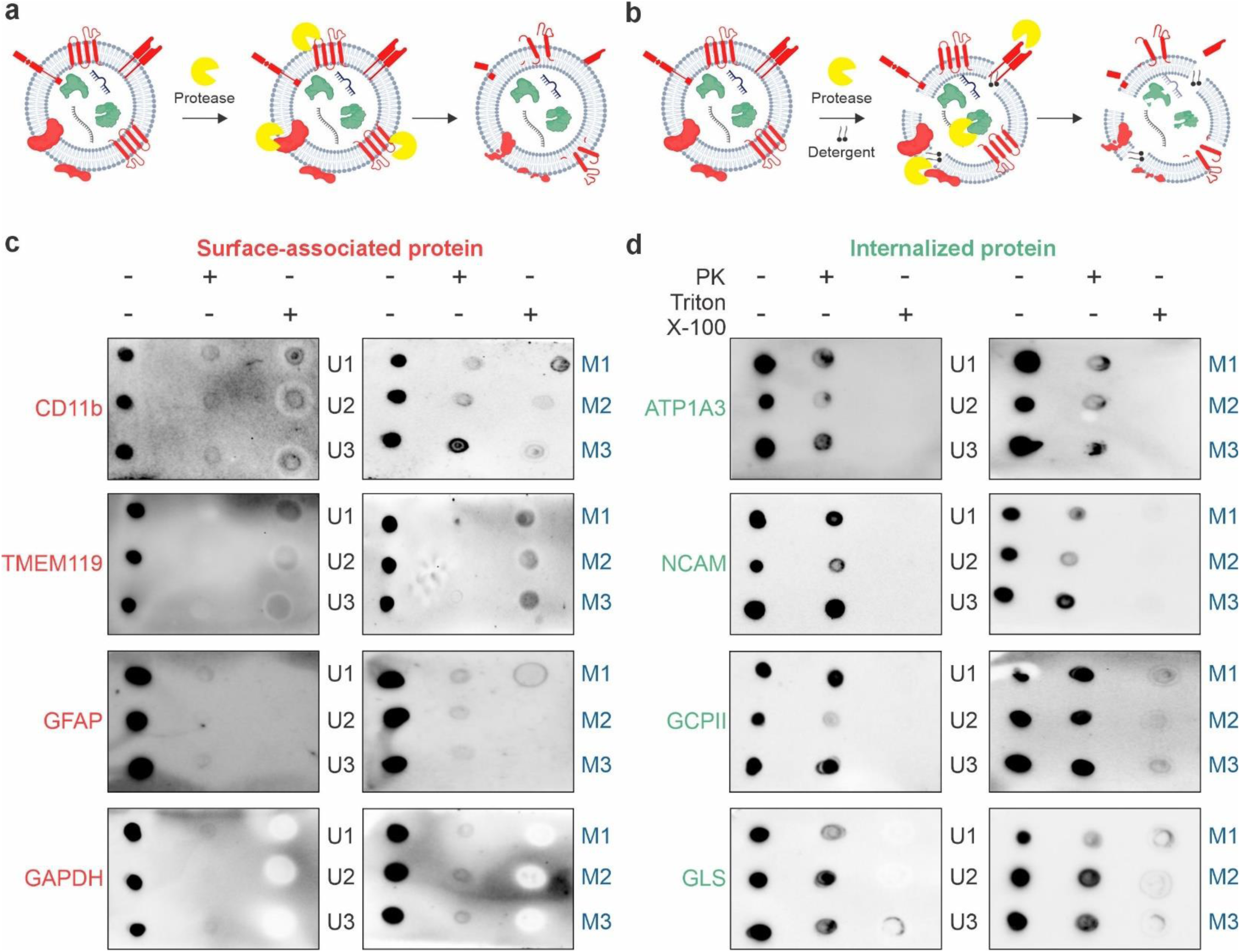
Proteinase K protection assay reveals differential localization of EV-associated proteins in brain-derived EVs. **a,b** Schematic representation of the proteinase K (PK) protection assay workflow. (**a**) Treatment of intact EVs with protease (i.e., PK) selectively digests surface-exposed proteins, while membrane-protected or luminal proteins remain intact. Schematic was created with BioRender and CorelDraw2022. (**b**) Co-treatment with detergent (i.e., Triton X-100) permeabilizes EV membranes, allowing PK to degrade all EV-associated proteins. PK activity was inhibited using phenylmethylsulfonyl fluoride (PMSF) prior to downstream analysis. **c,d** Dot blot analysis of EVs isolated from uninfected control (U#) and *M. tuberculosis*-infected (M#) rabbit brain tissue treated with either protease (+PK), detergent and protease (+PK +Triton X-100), or left untreated. Three biological replicates performed per group. (**c**) CD11b, TMEM119, GFAP, and GAPDH were digested by PK regardless of detergent treatment, indicating they are surface-associated EV proteins. (**d**) ATP1A3, NCAM, GCPII, and GLS were preserved in the +PK condition but lost upon detergent permeabilization, indicating they are internalized EV proteins. Schematic was created in BioRender and CorelDRAW 2022.

## DISCUSSION

EVs are increasingly recognized as active participants in infection, capable of distributing molecular cargo that can propagate inflammatory signaling and contribute to disease progression. This concept is well supported in pulmonary TB, where EVs from biofluids and immune cell cultures show infection-induced changes in bioactive cargo^56–60^. In TB meningitis, a proof-of-concept study demonstrated that urine EVs can carry *M. tuberculosis* DNA detectable by PCR^61^. Nonetheless, EV cargo alterations within the CNS in response to infection remain largely unexplored. To the best of our knowledge, this study provides the first evidence that *M. tuberculosis* infection alters EV protein cargo in the brain, revealing molecular signatures relevant to TB meningitis pathogenesis. Our findings underscore the potential of brain-derived EVs to serve as both biomarkers and active mediators of CNS pathology during *M. tuberculosis* infection.

Methodologically, we demonstrate that the “EV-release” workflow is feasible in rabbit brain tissue, extending recent tissue-based protocols developed for mouse and human brain to rabbit samples. This approach, which isolated EVs released *ex vivo* from intact brain pieces rather than by enzymatic dissociation, recovers small EV–enriched populations with preserved morphology and molecular profiles^24^. We show that this method is robust and yields EVs with comparable yield, size, and morphology from both fresh and flash-frozen tissue, aligning with prior reports^62,63^. The ability to use frozen tissue is particularly advantageous in BSL-3 environments, where immediate processing is often impractical, and broadens the scope of EV studies in CNS infection models.

Our studies show that TB meningitis drives quantitative and qualitative changes in brain-derived EVs. *M. tuberculosis* infection significantly increased EV biogenesis, although the size and morphology were similar to EVs isolated from uninfected brains. Importantly, our proteomic analysis revealed that proteins enriched in brain-derived EVs during infection were strongly associated with glial-related pathways, particularly those linked to microglial and astrocytic function, whereas proteins associated with neuronal processes were downregulated. These results align with our previous work showing robust microglial activation and increased density in our rabbit model^10,11^ and provides additional supportive data that activated glial cells are major contributors to pathology during infection. Several neuroinflammatory models have shown microglia and astrocytes activated by proinflammatory stimuli increase EV secretion^64–67^ and EVs proinflammatory effects are well-demonstrated by several in vitro and in vivo models (e.g., cerebral ischemia, Alzheimer’s disease, viral encephalitis)^15,16,68^. Although not TB-specific, those reports support the concept that increased brain EV secretion may reflect microglial and astrocytic activation in response to infection. By contrast, the relative depletion of neuron-associated proteins, including synaptic and axonal components, may reflect reduced neuronal EV secretion, impaired neuronal integrity, or increased neuronal injury. Similar patterns have been reported in Alzheimer’s disease and traumatic brain injury, where glial EVs dominate vesicle increase at the expense of neuronal contributions^69^. Collectively, these observations suggest that *M. tuberculosis* infection alters the cellular balance of EV production in the brain, with glial cells emerging as major contributors to the vesicle landscape.

Proteomic profiling of brain-derived EVs revealed that *M. tuberculosis* infection significantly alters EV cargo to promote immune activation and impair neuroprotection. EVs from infected brain were enriched in host immune and stress-response proteins, with upregulated proteins associated with antigen presentation, vesicle trafficking, interferon-driven responses, and, importantly, host response to TB. Infection-specific EV cargo showed that EVs selectively incorporate proteins associated with microglial activation and immune signaling such as interferon, inflammasome, and IL-17 signaling, as well as TB host defense, reflecting a potential role for brain-derived EVs in amplifying immune-activating signals during *M. tuberculosis* infection. Conversely, brain-derived EVs had reduced proteins linked to neuroprotection, mitochondrial function, and neurodevelopment, including cholesterol biosynthesis and axon guidance. We have previously shown in this rabbit model that there is a downregulation of cholesterol and several sterol intermediates and alterations in EV biogenesis within the brain may be one mechanism by which cholesterol homeostasis is affected^70^. Together, these findings support a dual role for EVs in TB meningitis where they promote neuroinflammatory signaling while simultaneously diminishing protective neuronal support, providing a mechanistic framework for the persistent neurological injury seen in survivors.

Comparative proteomics also shows a strong link between EVs, glial activation, glutamine metabolism and neuronal excitotoxicity in TB meningitis. We found that EVs from infected brains were enriched in GLS, the enzyme that metabolizes glutamine to glutamate. Neurological deficits in TB meningitis have been linked to glutamate excitotoxicity, with transcriptomics and metabolomic studies of patient plasma and CSF showing increased glutamate and enrichment of glutamate excitotoxicity and NMDA receptor binding pathways^8,9,71^. Although the causative pathways leading glutamate excitotoxicity in TB meningitis are not known, GLS expression is upregulated in *M. tuberculosis-*infected macrophages and patients with pulmonary TB^72,73^. GLS is also increased in many CNS infections (e.g., HIV, Japanese encephalitis virus)^74,75^ and in neuroinflammatory conditions (e.g., Rett syndrome, cerebral ischemia)^68,76^. Importantly, GLS is a key mediator of neuroinflammation, enhancing microglial activation, and directly regulates vesicle biogenesis^15,67,68^. When GLS is packaged in EVs, as our studies showed with *M. tuberculosis* infection, it establishes a proinflammatory microenvironment and leads to neurotoxicity^15,75,77^. Indeed, GLS inhibitors have been shown in several models to decreased microglial activation, EV production, neurotoxicity, and functional impairment^15,68,76–78^. Moreover, EV-borne GLS may reprogram recipient cell metabolism by promoting glutamate production and fueling the TCA cycle^78^. In the context of TB meningitis, GLS-enriched EVs could propagate inflammatory bioenergetic demands, promote excitotoxicity, and reinforce EV biogenesis pathways, providing a potential mechanistic explanation of its pathological consequences.

Bioinformatics predicts a useful starting point for inferring specific cell-type origins of EV populations. Protease protection assays further complemented this approach by distinguishing surface-exposed and internalized proteins, thereby identifying potential capture targets for cell-type EVs, such as microglia-derived EVs. In our study, TMEM119, CD11b, GFAP, and GAPDH were protease-sensitive, indicating surface exposure. Although GAPDH is cytosolic, its surface accessibility is compatible with prior reports of noncanonical surface display or corona adsorption on EVs^79^. Conversely, ATP1A3, NCAM, GCPII, and GLS were protease-resistant and localized internally. Prior studies identified NCAM1 and ATP1A3 on EV surfaces^80,81^, which differ from our results. These discrepancies likely reflect a higher fraction of endosome-derived small EVs relative to shedding microvesicles, species or brain-region context, infection-driven post-translational modifications that mask epitopes, or protein corona that shields or reorients antigens. Technical variables, including EV isolation methods, protease exposure (i.e., concentration, incubation time), and antibody epitope location may also contribute. Although CellMarker 2024 categorized GLS as a microglia-derived EV, our data indicates that other surface-associated proteins, such as TMEM119 or CD11b for microglia and GFAP for astrocytes, will be required to isolate cell-specific EVs to understand their individual contribution to infection pathogenesis.

Our study has several limitations. We isolated EVs from our experimentally-induced TB meningitis model that uses direct inoculation which does not fully recapitulate *M. tuberculosis* dissemination to the brain. We relied on protein enrichment to predict likely cellular origin rather than exclusive cell-type specificity, as several markers are shared across lineages. Protein localization assignments are based on protease-accessibility assays that assume intact membranes and minimal protein corona; these should be corroborated with orthogonal approaches such as immunoelectron microscopy, surface biotinylation with membrane-impermeant reagents, or surface proteomics. In addition, species differences must be considered, as EV markers validated in human brain may not directly translate to rabbits. Despite these limitations, our findings provide a strong foundation for future mechanistic and translational studies aimed at leveraging EVs as biomarkers and therapeutic targets in TB meningitis.

In summary, EVs from infected brains contained diverse proteome changes reflecting both immune activation and neuronal stress. The identification of proteins linked to TB pathogenesis, host immune regulation, and neurodegenerative processes underscores the potential of EVs as a window into CNS host–pathogen interactions. We provide experimental evidence that *M. tuberculosis* infection alters both composition and abundance of brain-derived EVs, underscoring their dual role as biomarkers and mediators of CNS pathology. Comparative proteomics revealed infection-specific signatures that represent molecular fingerprints of TB meningitis. These data demonstrate the feasibility and importance of EV profiling in CNS infection and set the stage to define their diagnostic or therapeutic utility. Future studies should refine cell-type–specific EV capture, validate EV marker accessibility across species and disease contexts, and determine whether these EV signatures can serve as biomarkers of disease severity, prognosis, or treatment response in TB meningitis. Single-vesicle technologies such as nanoFCM or single-particle interferometric imaging could quantify neuronal versus glial EV contributions and determine whether inflammatory cargo co-localizes with cell-type markers. Functional studies could also provide crucial information on how EVs from infected brain tissue influence neuronal survival, GLS expression and immune activation.

## ACKNOWLEDGEMENTS

This work was funded by the U.S. National Institutes of Health Allergy and Infectious Diseases grant K08-AI139371 (E.W.T.), Johns Hopkins Anesthesiology and Critical Care Medicine StAAR Investigator Award (E.W.T.), Johns Hopkins Children’s Center Innovation Grant (E.W.T.), and Johns Hopkins Tuberculosis Research Advancement Center (JHU TRAC) Development Award 5P30AI168436 (E.W.T.). D.B.F. was supported by U.S. National Institutes of Health National Heart, Lung, and Blood Institute grant R01HL164478 and U.S. Army Medical Research Grant HT94252410277. S.D. was supported by U.S. National Institutes of Health grant U18TR003780 and the American Heart Association grants TPA970850 and 23DIVSUP1057308. TEM was performed at the Johns Hopkins University School of Medicine Institute for Basic Biomedical Sciences Microscope Facility.

## AUTHOR CONTRIBUTIONS

B.V.R. and E.W.T. conceptualized and designed the studies. E.W.T., B.V.R., and N.N.L.D. performed studies in the rabbit model of TB meningitis. B.V.R. and N.N.L.D. performed EV isolation. B.V.R., N.N.L.D., N.B., and D.P. performed EV characterization. D.B.F. performed proteomic data analysis. E.W.T., B.V.R., N.N.L.D., and S.D. performed data analysis and interpretation. B.V.R. and E.W.T. wrote the original draft. All co-authors participated in the review and editing of the manuscript and approved the final version for submission.

## DATA AVAILABILITY

All data are available in the main text or the supplementary materials. The mass spectrometry proteomics data have been deposited to the ProteomeXchange Consortium via the PRIDE^82^ partner repository with the dataset identifier PXD071017 and 10.6019/PXD071017.

