## Supplemental Material for "Tuberculous meningitis alters the proteomic landscape of brain-derived extracellular vesicles"

Elizabeth W. Tucker

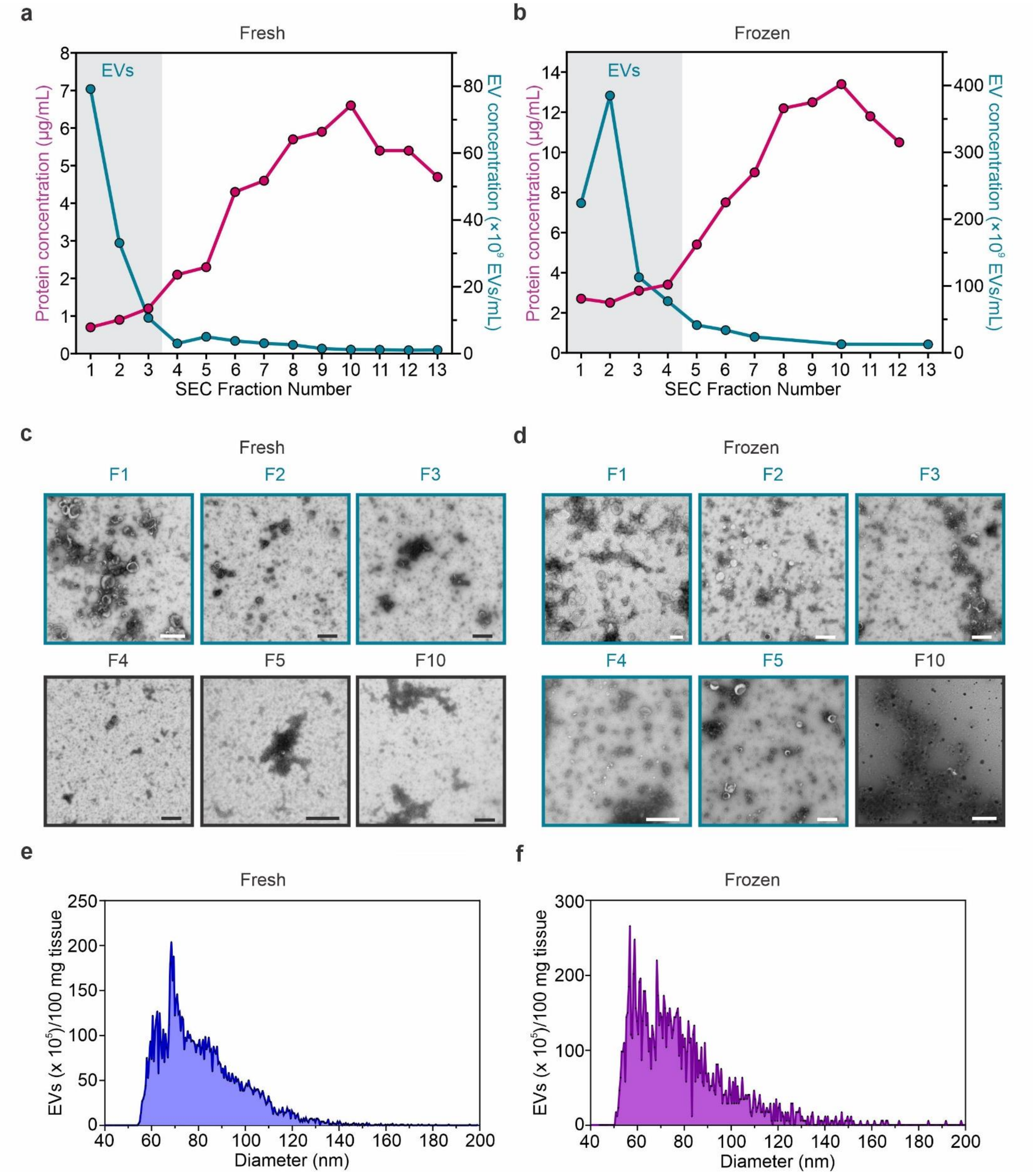


**Fig S1. Brain-derived EVs from healthy adult rabbits using fresh and flash-frozen tissue processing.** EVs were isolated from brain tissue from healthy adult rabbits for EV isolation protocol optimization. N = 1 rabbit/group. **a,c,e** Adult rabbit brain processed freshly at the time of euthanization. **b,d,f** Adult rabbit brain processed after flash freezing in -80°C. **a,b** Protein and EV concentration from first 13 size exclusion chromatography (SEC) fractions from the EV isolation from (**a**) freshly-processed and (**b**) flash-frozen adult rabbit brain. **c,d** Negative-stain TEM visualized brain-derived EVs from adult rabbit brain with teal boxes indicating EV-rich fractions, corresponding to fraction 1-3 (F1-F3) from (**c**) freshly-processed and F1-F5 from (**d**) flash-frozen brain tissue. Scale bar = 500 nm. **e,f** Size distribution determined by Nanoflow cytometry (NanoFCM) for (**e**)

freshly-processed and (**f**) flash-frozen adult rabbit brain. EV quantification was normalized per 100 mg of brain tissue.

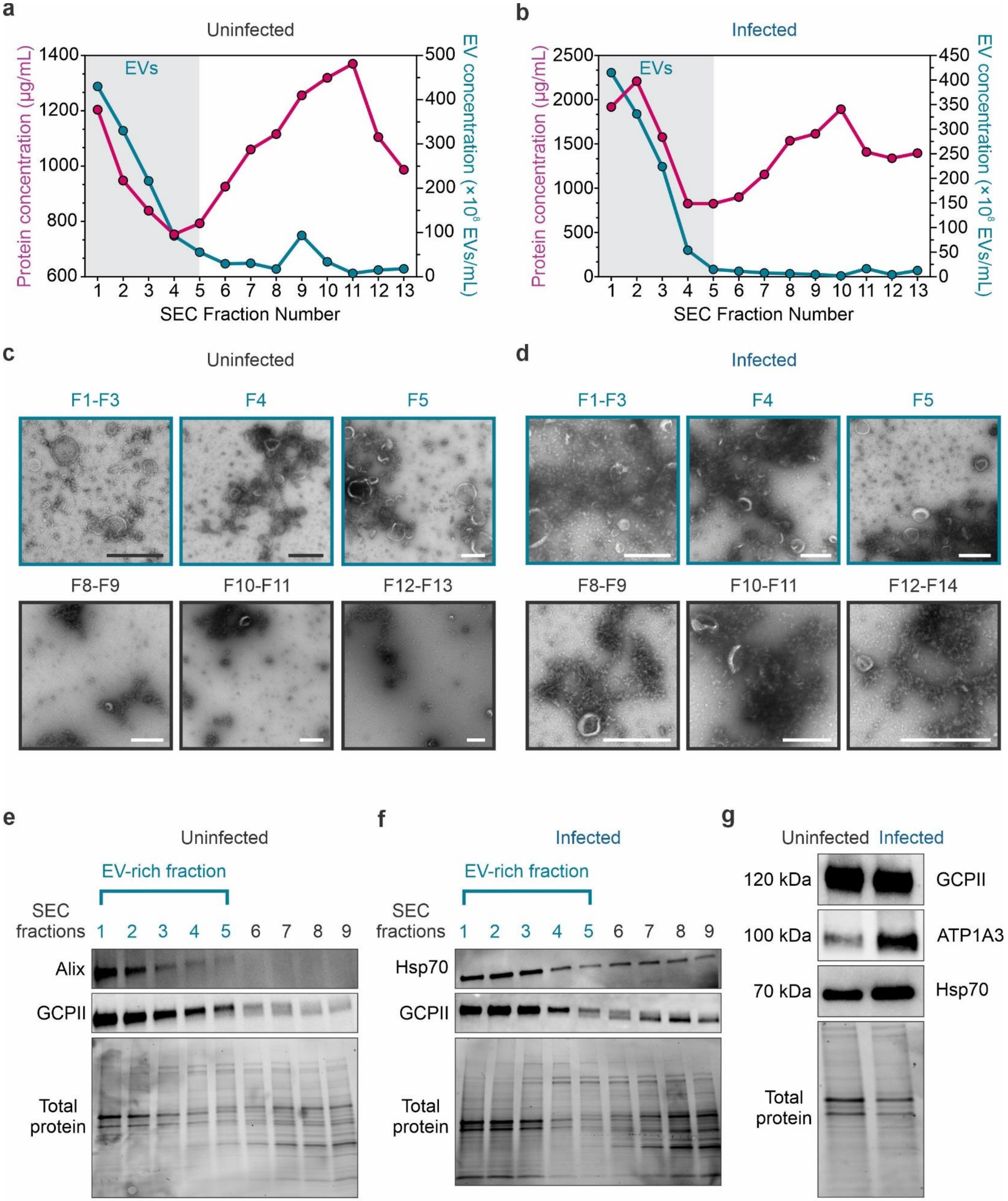


**Fig S2. Characterization of brain-derived EV isolation from 1-week old rabbits.** Brain-derived EVs were isolated from brain tissue from 1-week-old uninfected and infected rabbits (1-day post-injection of *M. tuberculosis*). N = 1 rabbit/group. **a,b** Protein concentration by BCA and EV concentration by NanoFCM from the first 13 size exclusion chromatography (SEC) fractions in (**a**) uninfected and (**b**) infected rabbits. EVs are enriched in F1-F5. **c,d** Negative-stain TEM visualized brain-derived EVs from several individual and pooled SEC fractions in (**c**) uninfected and (**d**) infected rabbits. Scale bar = 500 nm. **e,f** Western blots of common EV positive markers (Alix, Hsp70,) and the glial-associated protein, GCPII, in SEC fractions from (**e**) uninfected and (**f**) infected rabbits. Total protein loading is shown below each blot. In both groups, EV-associated markers were primarily detected in EV-rich fractions (F1–F5). **g** Pooled EV-rich fractions (F1–F5) from each condition were analyzed by Western blot for GCPII (glial-associated protein), ATP1A3 (neuronal protein), and Hsp70 (EV-positive marker). GCPII levels appeared comparable between groups, while ATP1A3 and Hsp70 were more abundant in EVs from the *M. tuberculosis*-infected brain. Total protein staining confirms equal protein loading across conditions.

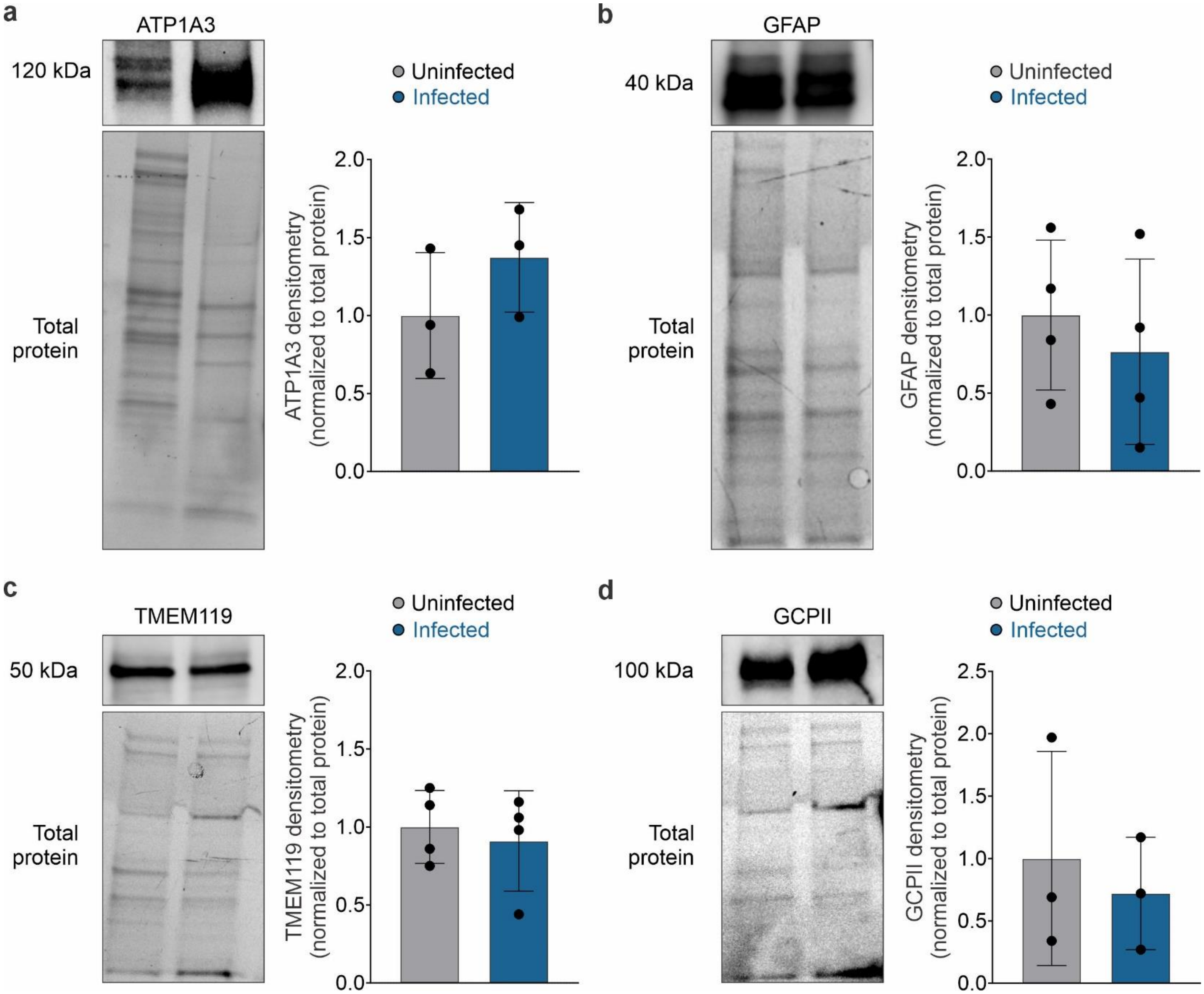


**Fig S3. Expression of cell type–associated proteins in brain-derived EVs. a-d** Western blot (left) and corresponding densitometry (right) of (**a**) ATP1A3 (neuronal), (**b**) GFAP (astrocytic), (**c**) TMEM119 and (**d**) GCPII (microglial/glial). All protein levels were normalized to total protein loading. Data are presented as mean ± SD from biologically independent samples. N = 3-4 rabbits/group. Statistical comparisons were made using unpaired two-tailed t-test.
